# Biocompatibility of Subcutaneously Implanted Plant-Derived Cellulose Biomaterials

**DOI:** 10.1101/039644

**Authors:** Daniel J. Modulevsky, Charles M. Cuerrier, Andrew E. Pelling

**Author notes:** These authors contributed equally to this work (Daniel J. Modulevsky, Charles M. Cuerrier). Author for correspondence: Andrew E. Pelling, 150 Louis Pasteur, MacDonald Hall, University of Ottawa, Ottawa, ON K1N 6N5, Canada, Web: http://www.pellinglab.net.

## Abstract

There is intense interest in developing novel biomaterials which support the invasion and proliferation of living cells for potential applications in tissue engineering and regenerative medicine. Decellularization of existing tissues have formed the basis of one major approach to producing 3D scaffolds for such purposes. In this study, we utilize the native hypanthium tissue of apples and a simple preparation methodology to create implantable cellulose scaffolds. To examine biocompatibility, scaffolds were subcutaneously implanted in wild-type, immunocompetent mice (males and females; 6-9 weeks old). Following the implantation, the scaffolds were resected at 1, 4 and 8 weeks and processed for histological analysis (H&E, Masson’s Trichrome, anti-CD31 and anti-CD45 antibodies). Histological analysis revealed a characteristic foreign body response to the scaffold 1 week post-implantation. However, the immune response was observed to gradually disappear by 8 weeks post-implantation. By 8 weeks, there was no immune response in the surrounding dermis tissue and active fibroblast migration within the cellulose scaffold was observed. This was concomitant with the deposition of a new collagen extracellular matrix. Furthermore, active blood vessel formation within the scaffold was observed throughout the period of study indicating the pro-angiogenic properties of the native scaffolds. Finally, while the scaffolds retain much of their original shape they do undergo a slow deformation over the 8-week length of the study. Taken together, our results demonstrate that native cellulose scaffolds are biocompatible and exhibit promising potential as a surgical biomaterial.

## INTRODUCTION

The development of novel biomaterials for tissue engineering strategies is currently under intense investigation [1–3]. Biomaterials are being developed for the local delivery of therapeutic cells to target tissues [4,5], the regeneration of damaged or diseased tissues [6–9] or the replacement of whole organs [10–15]. In their most general form, biomaterials provide a three-dimensional (3D) scaffold which attempts to mimic the *in vivo* cellular milieu [14,16]. Approaches have been developed to engineer the mechanical [17–24], structural [25] and biochemical properties [26–29] of these scaffolds with varying complexity. As well, significant efforts are underway to ensure that such implanted biomaterials are biocompatible and stimulate only minimal immune responses. The efforts in biomaterials research is being driven by the significant need for replacement organs and tissues. With an aging population, the gap between patients waiting for organ transplants and available donor organs is rapidly increasing [30]. While clinical applications of biomaterials have been somewhat limited, physicians have successfully utilized synthetic biomaterials to treat various damaged tissues and structures, such as skin, gum, cartilage, and bone [31–36].

Biomaterial scaffolds can take several forms such as powders, gels, membranes, and pastes [1,2]. Such polymer or hydrogel formulations can be moulded or 3D-printed to produce forms that are of therapeutic values [37–39]. An alternative approach to these synthetic strategies is whole organ decellularization [10,12–16]. Indeed, it has been shown that it is possible to dissociate the cells from a donated organ, leaving behind the naturally occurring scaffold matrix, commonly referred as a ghost organs [14]. The ghost organs lack any of the cells from the donor and can be subsequently cultured with cells derived from the patient or another source. Such approaches have already been utilized to repair and replace defective tissues [40–42]. In the past several years, many body parts have been created using synthetic and decellularization approaches, including the urethra, vaginal, ear, nose, heart, kidney, bladder, and neurological tissues [14,38,39,43–47].

However, these approaches are not without some disadvantages [48]. Synthetic techniques can require animal products and decellularization strategies still require donor tissues and organs. There has also been intense investigation into the development of resorbable biomaterials [49]. In these cases, the aim is to provide the body with a temporary 3D scaffold onto which healthy tissues can form. After several week or months, the implanted scaffold will be resorbed leaving behind a completely natural healthy tissue [26,29,50,51]. Although this is an ideal approach, many non-resorbable biomaterials (ceramic, titanium) have been successfully employed in clinical settings and play a major role in numerous therapies [2,49,52–57]. Importantly, resorbable biomaterials suffer from the fact that regenerated tissues often collapse and become deformed due to the loss of structure [58–62]. For example, for several decades, research on ear reconstruction from engineered cartilage has shown that biomaterial implants eventually collapse and become deformed as the implanted scaffolds break down and resorb [63]. However, recent successful approaches have relied on the use of resorbable collagen scaffolds embedded with permanent titanium wire supports [53,64,65]. Therefore, the need for non-resorbable, yet biocompatible, scaffolds persists in the field of tissue and organ engineering.

Recent complementary approaches have utilized scaffolding materials that are not derived from human organ donors or animal products. Namely, various forms of cellulose have been shown to have utility in both *in vitro* and *in vivo* studies [66–71]. Cellulose is abundant in nature, is easily produced and sourced, can be chemically modified to control surface biochemistry and produced as hydrogels with tuneable porosity and mechanical properties [67,72–77]. Moreover, nanocrystalline, nanofibrillar and bacterial cellulose constructs and hydrogels also have been shown to support the proliferation and invasion of mammalian cells *in vitro* and *in vivo* with high biocompatibility [78–83]. In our recent work, we developed an orthogonal, yet complementary, approach to organ decellularization and synthetic cellulose strategies. We developed a highly robust and cost effective strategy for producing cellulose biomaterials from decellularized apple hypanthium tissue [27]. The scaffolds required no further complex processing as is often the case in the production of nanocrystalline, nanofibrillar and bacterial cellulose constructs. The cellulose scaffolds were employed for *in vitro* 3D culture of NIH3T3 fibroblasts, mouse C2C12 muscle myoblasts and human HeLa epithelial cells. Our previous work revealed that these cells could adhere, invade and proliferate within the cellulose scaffolds and retain high viability even after 12 continuous weeks of culture.

Our previous work opens the question of *in vivo* biocompatibility [27]. Therefore, the objective of this study is to characterize the response of the body to apple-derived cellulose scaffolds. Macroscopic (~25 mm^3^) cell-free cellulose biomaterials were produced and subcutaneously implanted in mouse model for 1, 4 and 8 weeks. Here, we assess the immunological response of immunocompetent mice, deposition of extracellular matrix on the scaffolds and evidence of angiogenesis (vascularization) in the implanted cellulose biomaterials. Notably, although a foreign body response was observed immediately post-implantation, as expected for a surgical procedure, by the completion of the study only a low immunological response was observed with no fatalities or noticeable infections whatsoever in all animal groups. Surrounding cells were also found to invade the scaffold, mainly activated fibroblasts, and deposit a new extracellular matrix. As well, the scaffold itself was able to retain much of its original shape and structure over the 8-week study. Importantly, the scaffolds clearly had a pro-angiogenic effect, resulting in the growth of functional blood vessels throughout the implanted biomaterial. Taken together, our work demonstrates that we can easily produce 3D cellulose scaffolds that are biocompatible, becoming vascularized and integrated into surrounding healthy tissues.

## MATERIAL AND METHODS

### Animals

All experimental procedures were approved by the Animal Care and Use Committee of the University of Ottawa. Wild-type C57BL/10ScSnJ mice (males and females; 6-9 weeks old; n= 7 mice for each group) were purchased from The Jackson Laboratory (Bar Harbor, Maine, USA) and breed in our facilities. All animals were kept at constant room temperature (±22°C) and humidity (~52%). They were fed a normal chow diet and were kept under a controlled 12 hours light/dark cycle.

### Cellulose scaffold preparation

As described previously [27], McIntosh Red apples (Canada Fancy) were stored at 4°C in the dark for a maximum of two weeks. In order to prepare apple sections, the fruit was cut with a mandolin slicer to a uniform thickness of 1.14±0.08mm, measured with a Vernier caliper. Only the outer (hypanthium) tissue of the apple was used. Slices containing visible ovary-core tissue were not used. The slices were then cut parallel to the direction of the apple pedicel into squares segments of 5.14±0.21mm in length and with an area of 26.14±1.76mm^2^. Apple tissue was then decellularized by using a well-established protocol [14] for removing cellular material and DNA from tissue samples while leaving behind an intact and three-dimensional scaffold. Individual apple tissue samples were placed in sterilized 2.5ml microcentrifuge tubes and 2ml of 0.1% sodium dodecyl sulphate (SDS; Sigma-Aldrich) solution was added to each tube. Samples were shaken for 48 hours at 180 RPM at room temperature. The resultant cellulose scaffolds were then transferred into new sterile microcentrifuge tubes, washed and incubated for 12 hours in PBS (Sigma-Aldrich). To sterilize the cellulose scaffold, they were incubated in 70% ethanol for 1 hour and then washed 12 times with PBS. The samples were then maintained in PBS with 1% streptomycin/penicillin (HyClone) and 1% amphotericin B (Wisent, QC, Canada). At this point, the samples were immediately used or stored at 4°C for no more than 2 weeks.

### Cellulose implantation

The mice were anesthetized using 2% Isoflurane USP-PPC (Pharmaceutical partners of Canada, Richmond, ON, Canada) and their eyes protected by the application of ophthalmic liquid gel (Alco Canada In., ON, Canada). To prepare the surgery sites, mouse back hairs were shaved and the skins were cleaned and sterilized using ENDURE 400 Scrub-Stat4 Surgical Scrub (chlorhexidine gluconate, 4% solution; Ecolab Inc., Minnesota, USA) and Soluprep (2% w/v chlorhexidine and 70% v/v isopropyl alcohol; 3M Canada, London, ON, Canada). To maintained animal hydration, 1ml of 0.9% sodium chloride solution was administrated subcutaneously (s.c.) (Hospira, Montréal, QC, Canada). During the surgical procedures, we applied all sterility measures requested for survival surgeries. To implant the scaffolds, two 8mm incisions were made on the dorsal section of each mouse (upper and lower). Two cellulose scaffold samples were separately and independently implanted on each mouse. The incisions were then sutured using Surgipro II monofilament polypropylene 6-0 (Covidien, Massachusetts, USA) and transdermal bupivicaine 2% (as monohydrate; Chiron Compounding Pharmacy Inc., Guelph, ON, Canada) was topically applied on surgery sites to prevent infection. Also, buprenorphine (as HCL) (0.03mg/ml; Chiron Compounding Pharmacy Inc. Guelph, ON, Canada) was administrated s.c. as a pain reliever. All animals were then carefully monitored for the next 3 days by animal care services and received repetitions of the same pharmacological treatments.

### Scaffold resections

At 1, 4 and 8 weeks after scaffold implantation, the mice were euthanized using CO_2_ inhalation. After blood collection, the dorsal skin was carefully resected and immediately immersed in PBS solution. The skin sections containing cellulose scaffolds were then photographed, cut and fixed in 10% formalin for at least 48 hours. The samples were then kept in 70% ethanol before being embedded in paraffin by the PALM Histology Core Facility of the University of Ottawa.

### Histological analysis

Serial 5μm thick sections were cut, beginning at 1 mm inside the cellulose scaffold, and stained with hematoxylin and eosin (H&E) and Masson’s trichrome. For immunocytochemistry, heat induced epitope retrieval was performed at 110°C for 12 min with citrate buffer (pH 6.0). Anti-CD31/PECAM1 (1:100; Novus Biologicals, NB100-2284, Oakville, ON, Canada), anti-alpha smooth muscle actin (1:1000, ab5694, abcam, Toronto, ON, Canada) and anti-CD45 (1:3000; ab10558, abcam, Toronto, ON, Canada) primary antibodies were incubated for an hour at room temperature. Blocking reagent (Background Sniper, Biocare, Medical, Concorde, CA, USA) and detection system MACH 4 (Biocare Medical, Concord, CA, USA) were applied according to company specifications. For the evaluation of cell infiltration, extracellular matrix deposition and vascularisation (angiogenesis), micrographs were captured using Zeiss MIRAX MIDI Slide Scanner (Zeiss, Toronto, Canada) equipped with 40x objective and analysed using Pannoramic Viewer (3DHISTECH Ltd., Budapest, Hungary) and ImageJ software. The scoring of inflammation was evaluated by a pathologist. The scoring was subjectively assigned by qualitative analysis of the magnitude of the total foreign response as well, the cell population proportions within the foreign response.

### Scanning electron microscopy (SEM)

The structure of cellulose was studied using a scanning electron microscopy. Globally, scaffolds were dehydrated through successive gradients of ethanol (50%, 70%, 95% and 100%). Samples were then gold-coated at a current of 15mA for 3 minutes with a Hitachi E-1010 ion sputter device. SEM imaging was conducted at voltages ranging from 2.00- 10.0 kV on a JSM-7500F Field Emission SEM (JEOL, Peabody, MA, USA).

### Statistical analysis

All values reported here are the average ± standard deviations. Statistical analyses were performed with one-way ANOVA by using SigmaStat 3.5 software (Dundas Software Ltd, Germany). A value of *p* < 0.05 was considered statistically significant.

## RESULTS

### Scaffold Preparation

Cellulose scaffolds were prepared from apple tissue using a modified decellularization technique we have previously described [27]. All scaffolds were cut to a size of 5.14±0.21 x 5.14±0.21 x 1.14±0.08mm (**Fig 1A**), decellularized and prepared for implantation (**Fig 1B**). The scaffolds appear translucent after decellularization due to the loss of all plant cellular material and debris. The removal of apple cells was also confirmed with histological observation (**Fig 1C**) and scanning electron microscopy (**Fig 1D**). Analysis of the histological images and the measurement of the average wall thickness (4.04±1.4μm) reveal that the cellulose scaffolds is highly porous, capable of being easily invaded by nearby cells and results in an acellular cellulose scaffold that maintains its shape.

**Figure 1:**
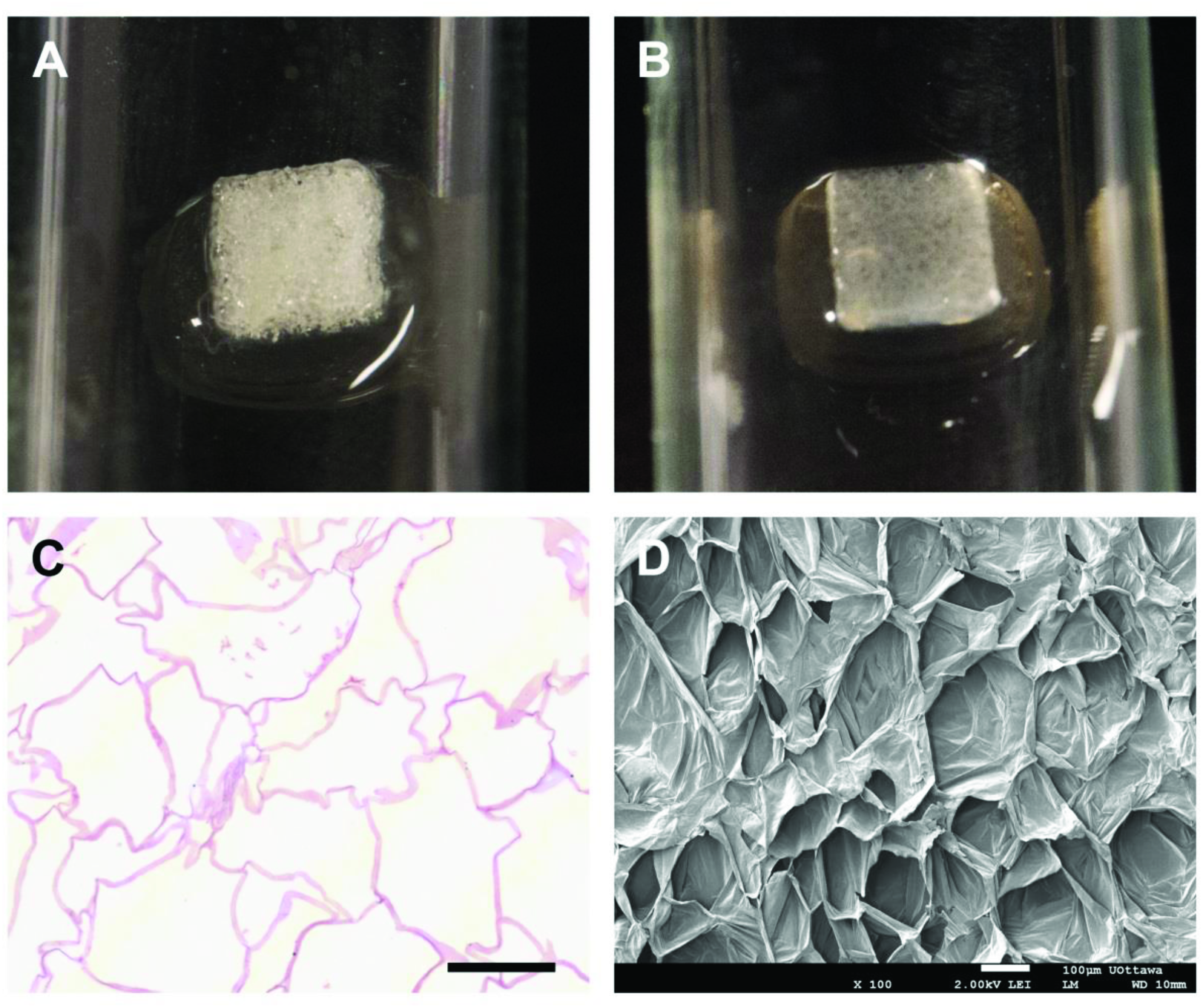
Cellulose scaffold preparation. Macroscopic appearance of a freshly cut apple hypanthium tissue (**A**) and the translucent cellulose scaffold biomaterial post-decellularization and absent of all native apple cells or cell debris (**B**). H&E staining of cross sectioned decellularized cellulose scaffold (**C**). The cell walls thickness and the absence of native apple cells following decellularization are shown. The 3D acellular and highly porous cellulose scaffold architecture is clearly revealed by scanning electron microscopy (**D**). Scale bar: **A-B** = 2mm, **C-D** = l00μm.

### Implantation of Cellulose Scaffolds

Two independent skin incisions (8mm) were produced on the back of each mouse to create small pouches for the biomaterial implantation (**Fig 2A**). One cellulose scaffold (**Fig 2B**) was implanted in each subcutaneous pouch. Throughout the study, there were no cases of mice exhibiting any pain behaviour that may have been induced by the cellulose scaffold implantation and none of them have displaying any symptoms of visible inflammation or infection. The cellulose scaffolds were resected at 1 week, 4 weeks and 8 weeks after their implantation and were photographed to measure the change in scaffold dimensions (**Fig 2D-F**). At all-time points, healthy tissue can be observed surrounding the cellulose scaffold with the presence of blood vessels, that are proximal or in direct contact, and the scaffolds retain their square shape. The pre-implantation scaffold had an area of 26.3±1.98mm^2^ and it was observed to slowly decrease as function of their implantation time base on the scaffold area that is visible to the naked eye on the skin (**Fig 2G**). At 8 weeks post-implantation, the scaffold dimensions reach a near plateau measurement of 13.82±3.88mm^2^ demonstrating an approximate 12mm^2^ (48%) change over the course of this study.

**Figure 2:**
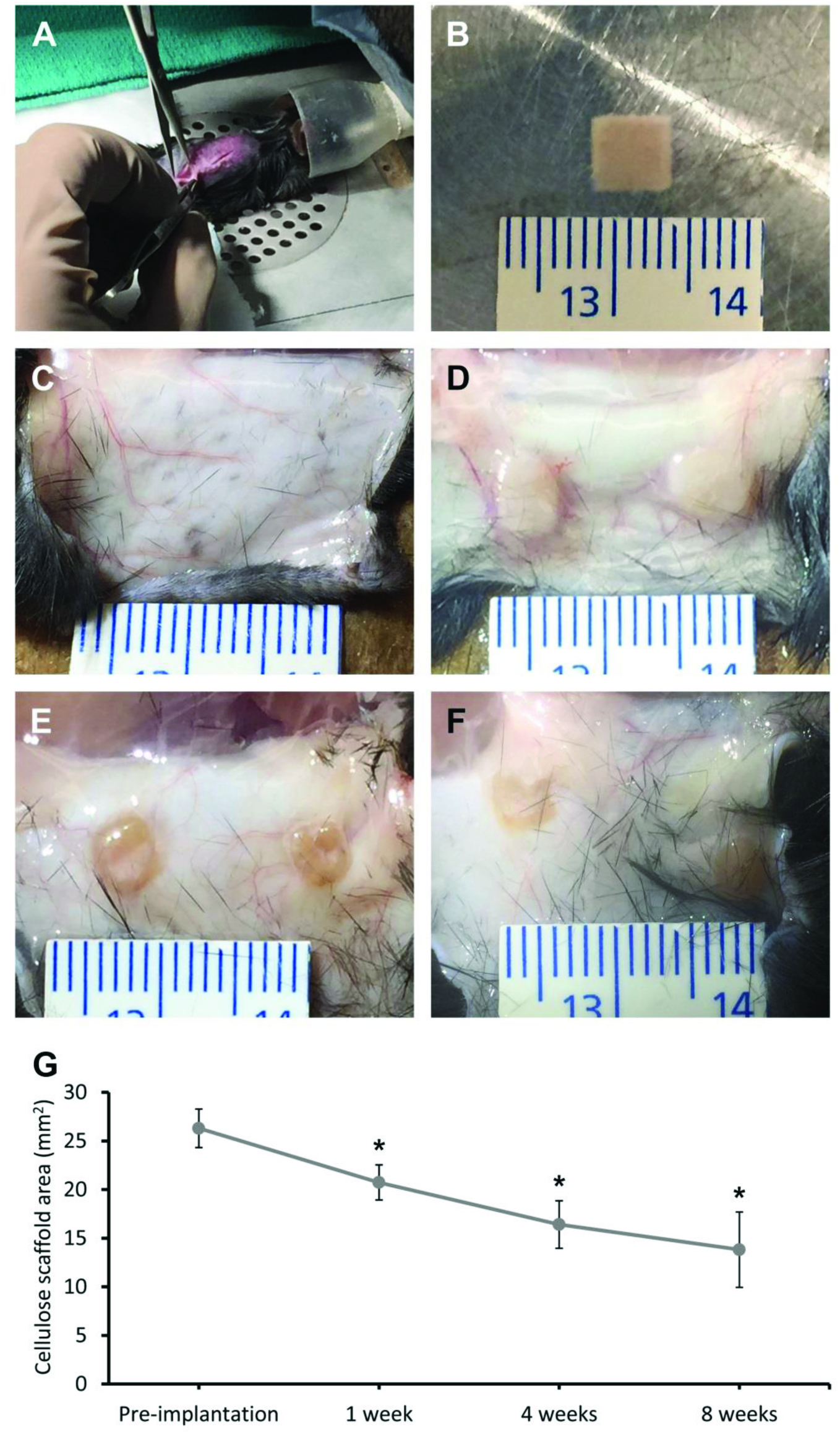
Cellulose scaffolds implantation and resection. The subcutaneous implantations of cellulose scaffolds biomaterial were performed on the dorsal region of a C57BL/10ScSnJ mouse model by small skin incisions (8 mm) (**A**). Each implant was measured before their implantation for scaffold area comparison (**B**). Cellulose scaffolds were resected at 1 week (**D**), 4 weeks (**E**) and 8 weeks (**F**) after the surgeries and macroscopic pictures were taken (control skin in **C**). The changes in cellulose scaffold surface area over time are presented (**G**). The pre-implantation scaffold had an area of 26.30±1.98mm^2^. Following the implantation, the area of the scaffold declined to 20.74±1.80mm^2^ after 1 week, 16.41±2.44mm^2^ after 4 weeks and 13.82±3.88mm^2^ after 8 weeks. The surface area of the cellulose scaffold has a significant decrease of about 12mm^2^ (48%) after 8 weeks implantation (* = *P*<0.001; n= 12-14).

### Biocompatibility and cell infiltration in plant derived cellulose scaffolds

Scaffold biocompatibility and cell infiltration was examined with H&E staining of fixed cellulose scaffolds at 1, 4 and 8 weeks following their implantation (**Fig 3**). The global views of longitudinal section of representative cellulose scaffolds are shown in **Fig 3A-C**. The scaffolds are implanted under the muscular layer of the dermis. Interstitial fluids, stained in pink, can be seen throughout the implanted scaffold, in contrast to a non-implanted scaffold (see **Fig 1C**), highlighting their high porosity and permeability. Within the global view it was observed that the scaffold maintains its general shape throughout the study. In **Fig 3D-F**, a magnified section of the perimeter of the scaffold is shown at each post-implantation time points. At 1 week, the dermis tissue surrounding implant displays symptoms of an acute moderate to severe immune response (qualitative study performed by a pathologist) (**Fig 3D**). As well a dense layer of cells can be seen infiltrating into the cellulose scaffolds. The population of cells within the scaffold at 1 week consist mainly of granulocytes, specifically; polymorphonuclear (PMN) and eosinophils (**Fig 3D**). There is also a population of dead cells and apparent cell debris. Importantly, all of these observations are completely consistent with an expected acute foreign body reaction that follows implantation [84–86]. At the 4 week point we observed a stark difference in both the surrounding epidermis tissue and in the cell population migrating into the cellulose scaffold (**Fig 3E**). The epidermis tissue surrounding the cellulose scaffold has a decreased immune response, now scored as mild to low. The population of cells within the epidermis surrounding scaffolds now contain higher levels of macrophages and lymphocytes (**Fig 3E**). This is an anticipated characteristic of the foreign body reaction to an implanted biomaterial, demonstrating the scaffold cleaning process [84–86]. There is also an increase in the population of multinucleated cells within the interior of the scaffold as part of an inflammatory response (**Fig 3E**). Finally, 8 weeks post-implantation, the immune response apparent at 1 and 4 weeks has completely disappeared (**Fig 3F**), with the epidermis tissue now appearing normal. In fact, the epidermis tissue in contact with the cellulose scaffold contains the same structures as normal epidermis tissue. In the cellulose scaffold perimeter there is now a lower density of cells due to the decreased inflammation and notably, there are no fragmented dead cells present. Instead, the population of cells now contain an elevated level of macrophages, multinucleated cells and active fibroblasts. The active fibroblasts (appearing spindle shaped), can be observed migrating from the surrounding epidermis into the cellulose scaffold. In fact, fibroblasts were found throughout the cellulose scaffold. These results demonstrate that by 8 weeks post-implantation the cellulose scaffold has been accepted by the host. In parallel with the H&E inflammation analysis, we performed anti-CD45 staining to evaluate the level of inflammation throughout the scaffold and surrounding dermis tissue (**Fig 3 G-I**). It is clear that the inflammation throughout the dermis and within the scaffold is elevated after 1 week. However, the amount of leukocytes significantly decreases in the surrounding dermis and scaffold over the implantation time reaching a near basal level at 8 weeks.

**Figure 3:**
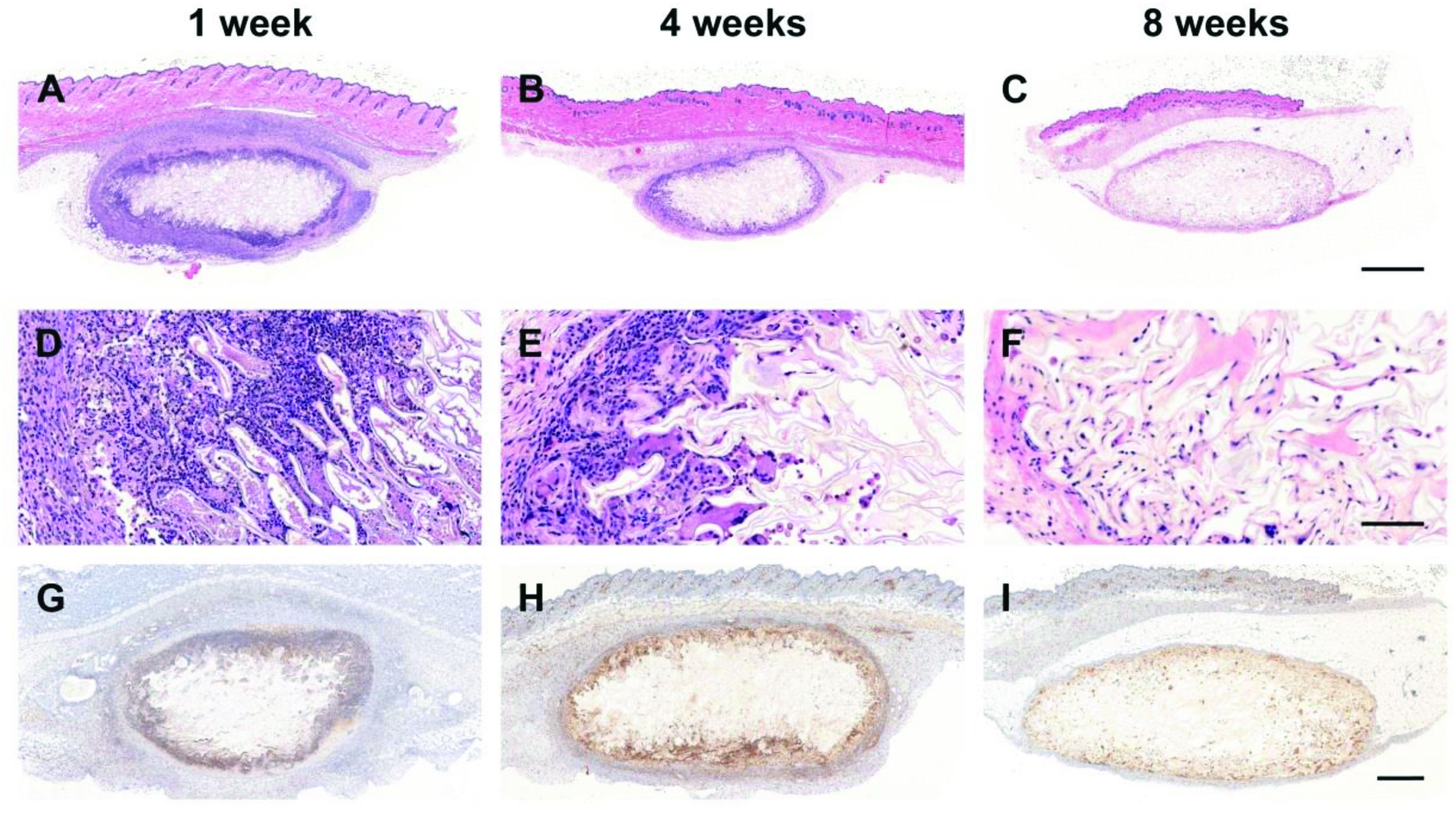
Biocompatibility and cell infiltration. Cross sections of representative cellulose scaffolds stained with H&E and anti-CD45. These global view show the acute moderate-severe anticipated foreign body reaction at 1 week (**A**), the mild chronic immune and subsequent cleaning processes at 4 weeks (**B**) and finally, the cellulose scaffold assimilated into the native mouse tissue at 8 weeks (**C**). Higher magnification regions of interest (**D-F**) allow the observation of all the cell type population within biomaterial assimilation processes. At 1 week, we can observe populations of granulocytes, specifically; polymorphonuclear (PMN) and eosinophils that characterize the acute moderate to severe immune response, a normal reaction to implantation procedures (**D**). At 4 weeks, a decreased immune response can be observed (mild to low immune response) and the population of cells within the epidermis surrounding scaffolds now contain higher levels of monocytes and lymphocytes characterizing chronic response (**E**). Finally, at 8 weeks, the immune response has completely resorbed with the epidermis tissue now appearing normal. The immune response observed with H&E staining is confirmed using anti-CD45 antibody, a well known markers of leukocytes (**G-I**). The population of cells within the scaffold are now mainly macrophages, multinucleated cells and active fibroblasts. Scale bars: **A-C** = 1mm, **D-F** = 100μm and **G-I** = 500μm.

### Extracellular Matrix Deposition in the Cellulose Scaffolds

The presence of active fibroblasts led us to question if the cellulose scaffold was acting as a substrate for the deposition of new extracellular matrix. This was determined using Masson’s Trichrome staining of fixed cellulose scaffolds slides at each time point following implantation (**Fig 4**). At 1 week post-implantation, the histological study shows the absence of collagen structures inside the collagen scaffold (**Fig 4A, D, and G**). As fibroblast cells invade the scaffold, as seen with H&E staining and confirmed by anti-alpha smooth muscle actin staining (data not shown), collagen deposits inside the cellulose scaffold can be observed after 4 weeks (**Fig 4B, E, and H**). At 8 weeks (**Fig 4C, F and I**) the collagen network is clearly visible inside the cavities of the cellulose scaffold. The complexity of the deposited collagen network is highlighted in **Fig 4I**, where we can detect individual collagen fibers within the collagen matrix. This is in contrast to the characteristic high density, thick, cable-like organization of collagen found in scar tissue.

**Figure 4:**
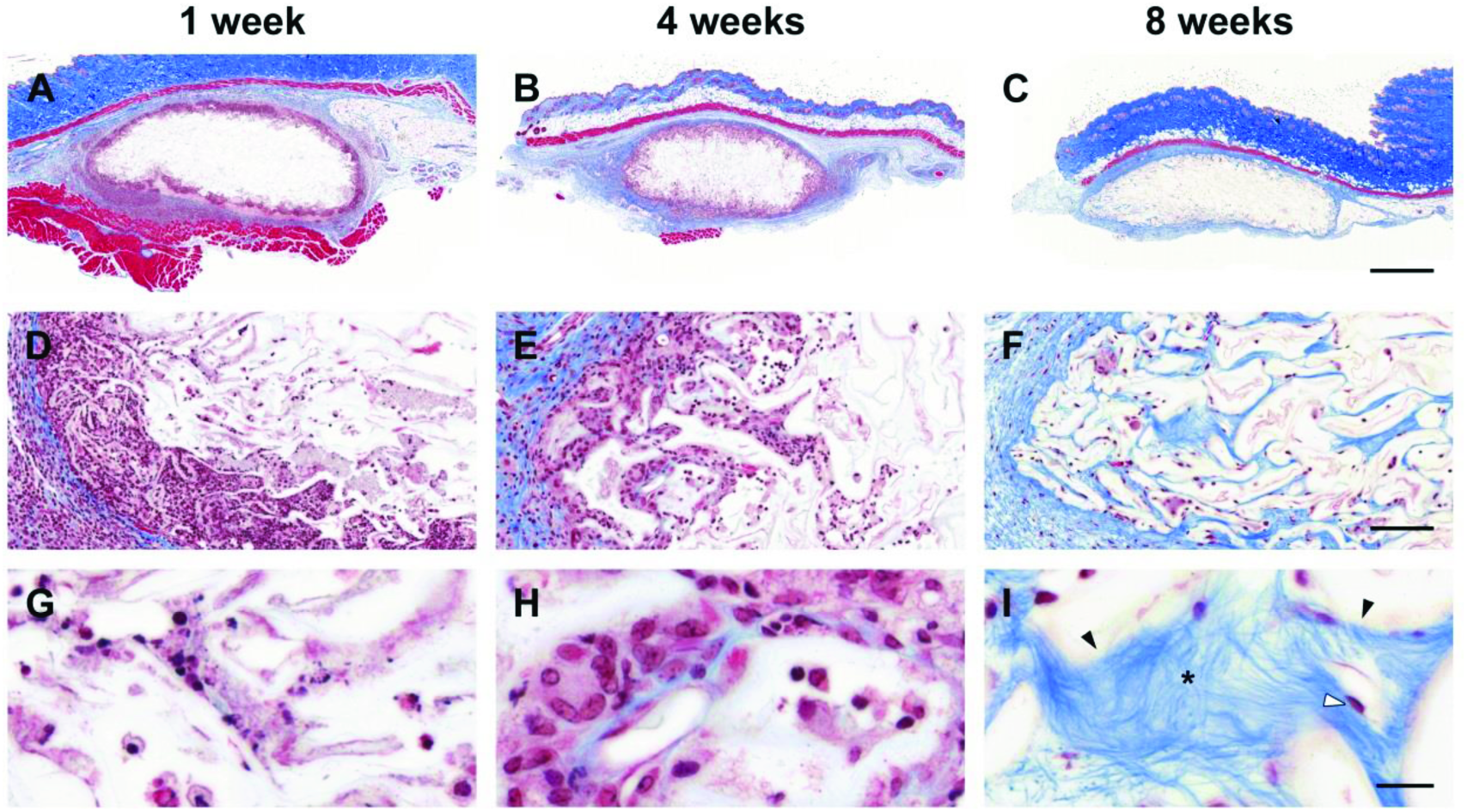
Extracellular matrix deposition. Cross sections of representative cellulose scaffolds stained with Masson’s Trichrome (**A-C**). After 1 week post-implantation, the magnification of region of interest in (**A**) show the lack of collagen structures inside the collagen scaffold (**D, G**). As fibroblast cells start to invade the scaffold, collagen deposits inside the cellulose scaffold can be sparsely observed after 4 weeks (**E, H**). Concomitant with the observation of activated fibroblast (spindle shaped cells) inside the cellulose scaffold, collagen network is clearly visible inside the cavities after 8 weeks (**F, I**). Scale bars: **A-C** = 1mm, **D-F** = 100μm and **G-I** = 20μm. * = collagen fibers; black arrows = cellulose cell wall; white arrow = fibroblast.

**Figure 5:**
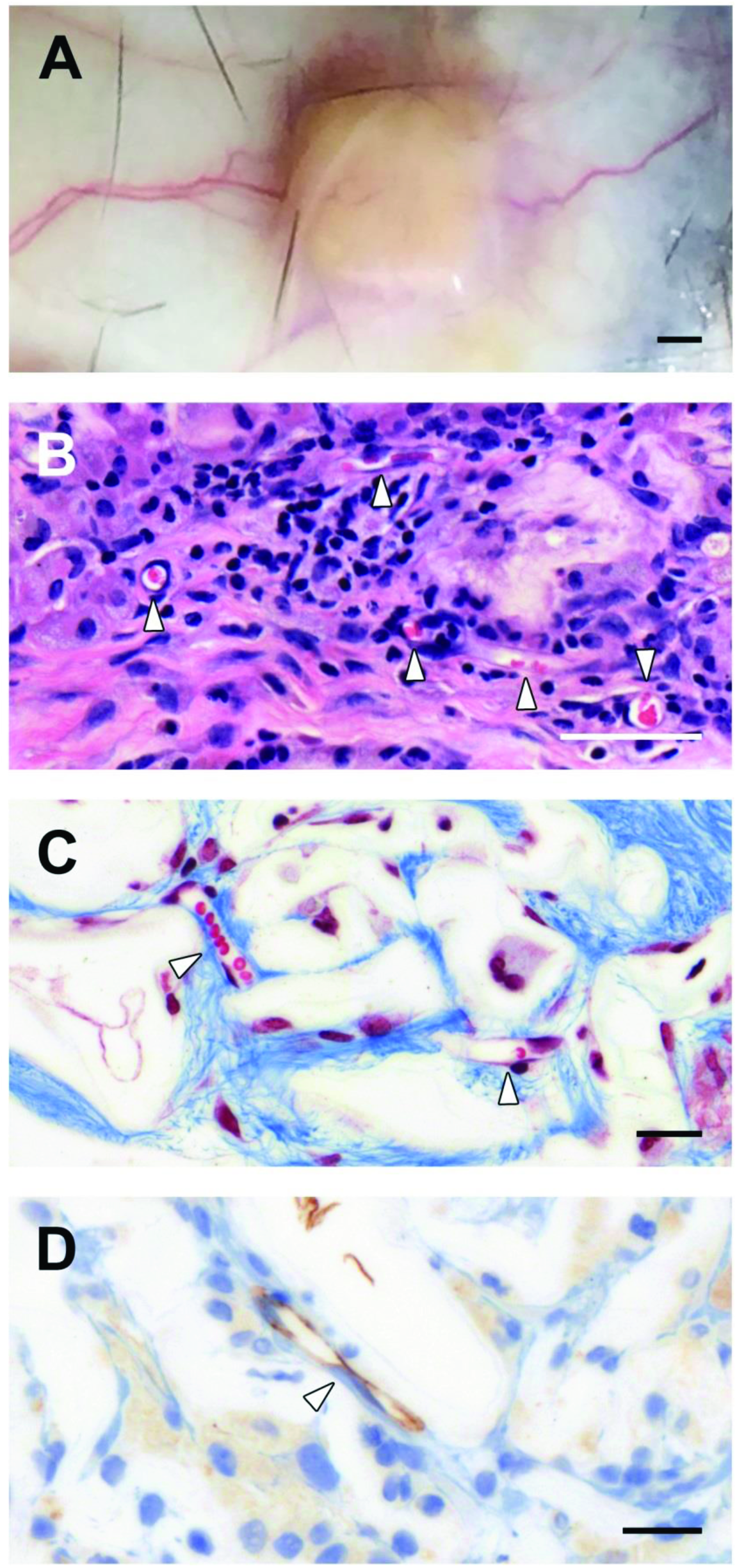
Vascularization and Angiogenesis. Macroscopic observations of blood vessels directly in the surrounding tissues around the cellulose scaffold (**A**). Confirmation of angiogenesis within the cellulose scaffold by the observation of multiple blood vessel cross sections in H&E staining (**B**) and Masson’s Trichrome staining (**C**) micrographs. The angiogenesis process was also confirmed with anti-CD31 staining to identify endothelial cells within the cellulose scaffold (**D**). Scale bars: **A** = 1mm, **B** = 50μm and **C-D** = 20μm. White arrows = blood vessels.

### Vascularization of the Cellulose Scaffolds

Capillaries ranging from 8 to 25μm were also identified within the scaffolds as early as 1 week post-implantation. At 4 week and 8 week post implantation, blood vessels and capillaries can be observed extensively within the scaffold and the surrounding dermal tissue. We observed blood vessels presence on the cellulose scaffold and in surrounding dermis in the macroscopic photos taken during the resection (**Fig 5A**). Multiple cross sections of blood vessels, with the presence of red blood cells (RBCs), are identified within 4 weeks of scaffold implantations (**Fig 5B**; H&E stain). The same results are obtained 8 weeks after implantation where capillaries with RBC and endothelial cells are clearly seen (**Fig 5C;** Masson’s Trichrome). All results on blood vessels formation were also confirmed with anti-CD31 staining to identify endothelial cells in the scaffold (**Fig 5D**).

## DISCUSSION

In this study, our objective was to examine the *in vivo* biocompatibility of acellular cellulose scaffolds derived from apple hypanthium tissue. To this end, acellular cellulose scaffolds were subcutaneously implanted within immunocompetent mice to establish their biocompatibility. Our data reveals that the implanted scaffolds demonstrate a low inflammatory response, promote cell invasion and extracellular matrix deposition, and act as a pro-angiogenic environment. Remarkably, none of the mice in this study died or demonstrated any symptoms of implant rejection such as edema, exudates or discomfort during the course of this research indicative of a successful implantation of the cellulose scaffolds. This implanted scaffolds are composed of a porous network of cavities in which the original host plant cells resided [69]. This architecture efficiently facilitates transfer of nutrients throughout the plant tissue. As we have shown here and in our previous study, the apple tissues are easily decellularized [27]. This simple treatment changes the appearance of the hypanthium tissue so that it becomes transparent, as a result of the removal of cellular materials.

Several important conclusions emerge from the current study. First, after implantation, the scaffolds are rapidly infiltrated with host cells, which begin with inflammatory cells. Consistent with previous findings, the immune response of the host animals followed a well-known timeline [84–88], ultimately demonstrating biocompatibility. As expected, the cell population within the scaffold after 1week post-implantation are mainly granulocytes, specifically; polymorphonuclear (PMN) and eosinophils, constituting a clear inflammatory response. The production of a provisional matrix around the scaffold was also observed resulting in an inflamed appearance in the tissue surrounding the scaffold [84–88]. This is not unexpected and is the result of the foreign material as well as a response to the surgical procedure [84–88]. Four weeks post implantation, the population of cells within the scaffold have evolved and are now lymphocytes, monocytes, macrophages, foreign body multinucleated cells as well as scattered eosinophils. Typical with chronic inflammation, the cellular debris present in the provisional matrix at 1 week, is now being cleared by the host immune system [84–88]. At 8 weeks, the cellulose scaffold is now void of all provisional matrix and cellular debris and low levels of macrophages and foreign body multinucleated cells are still visible within the scaffold. Consistent with the immune response within the cellulose scaffold, the surrounding tissue is observed to return to its original physiology. In fact, at 8 week implantation the surrounding tissue is nearly similar to control tissue. Although the immune response and inflammation at 8 weeks is low, low levels of macrophages can be observed within the scaffold. Although traditionally associated with inflammation, macrophages have beneficial roles consistent with our findings. Specifcially, macrophages are also known to secrete growth and pro-angiogenic factors, ECM proteins and pro-fibrogenic factors that actively regulate the fibro-proliferation and angiogenesis in tissue repair and regeneration [86]. Regardless, the vast population of cells within the scaffold after 8 weeks are now reactive fibroblasts. These cells are altering the microenvironment of the scaffold through the secretion of a new collagen extracellular matrix. Importantly the new matrix displays a remarkably low density compared, suggestive of regeneration as opposed to the characteristic high density, cable-like organization of collagen found in scar tissues [89].

Our data also demonstrates that the scaffolds are pro-angiogenic, which is critical to ensuring blood transport from the surrounding tissue [90]. As with native tissue, limited blood supply to the scaffold will result in ischemia and potentially necrosis. Interestingly, it was demonstrated that bioceramics with pore diameters lower than 400μm resulted in a decrease in the growth of blood vessels and limits the size of blood vessel diameter in *in vivo* implantations. The porous structure of the cell wall architecture is composed of overlapping cell wall cavities with diameters ranging from 100-300μm with manual interconnection distance of 4.04±1.4μm. As such, the high porosity size and low volume-fraction of the cellulose scaffolds are consistent with the promotion of blood vessel formation. Taken together, the cellulose scaffold now appears to completely void of the provisional matrix and fully accepted as a subcutaneous implant.

We also observed a decrease in the scaffold area over time, but it does not appear that the cellulose scaffold is in the processes of degradation. Rather, the change in area is due to the collapse of the cell wall cavities on the perimeter of the scaffold resulting from the active movement of the mouse. Active biological degradation is not expected to be possible as mammals lack the appropriate enzymes to digest plant-synthesized cellulose [91,92]. Moreover, the highly crystalline form of cellulose that is found in plant tissues is also known to resistant to degradation in mammals [92]. Alternatively, it has been demonstrated that *in vivo* cellulose implants can be chemically activated in order to be more easily degraded [93]. Most importantly however, highly crystalline forms of cellulose have some of the lowest reported immunological responses [92].

A large variety of clinically approved biomaterials are used to treat specific conditions within patients[1]. Such biomaterials can be derived from human and animal tissues, synthetic polymers, as well as materials such as titanium and ceramics [1,2,26,49,50,53,54,56,74,76,94–106]. However, these approaches are not without disadvantages that arise from concerns about the source, production costs and/or widespread availability [48]. There is currently an intense interest in developing resorbable biomaterials that will degrade *in vivo* and only act as a temporary scaffold that will promote and support the repair or regeneration of damaged/diseased tissue [49]. Although this is an ideal scenario, newly formed structures are also found to collapse as the scaffold degrade [53,64,107–109]. Moreover, the products of degradation can also be found to have toxic or undesirable side-effects [53,110,111]. For example, the reconstruction of the ear has become a well-known challenge in tissue engineering. Early studies have employed scaffolds in the shape of an ear that are produced from animal or human derived cartilage [53,58,59,61,63,64]. However, after implantation and eventual scaffold degradation, the ear is often found to collapse or deform [60–62]. Recent strategies have now opted to create biological composite materials composed of both a titanium frame embedded in a biological matrix [53]. Therefore, there is still a clear need for non-resorbable, yet inert and biocompatible, scaffolds persists in the field of tissue and organ engineering.

We suggest that plant-derived cellulose biomaterials offer one potential approach for the production of implantable scaffolds. This approach is complementary to bacterial cellulose strategies which have demonstrated clear utility as well [66,69–71,73,80,83,102,106,112–115]. However, plant derived materials are cost effective to produce and are extremely straightforward to prepare for implantation, exhibit clear biocompatibility, an ability to retain their shape while supporting the production of natural extracellular matrix and most importantly, the promotion of vascularization. In our previous work we have shown that the scaffolds can also be functionalized with proteins prior to culture *in vitro*. Such work will also be conducted in the future in order to explore the use of scaffold surface functionalization with growth factors and matrix proteins to promote the invasion of specific cell types, further minimize the early immune response and promote maximal vascularization. Moreover, the cellulose scaffolds can easily be formed into specific shapes and sizes, offering an opportunity to create new tissues with specific geometrical properties. Although there are numerous new avenues of research to follow, we have been able to demonstrate that acellular cellulose scaffolds are biocompatible *in vivo* in immunocompetent mice and might be considered as a new strategy for tissue regeneration.

## ACKNOWLEDGMENTS

We thank Joshua Lavigne, VT, RLAT for technical assistance with animals, Ana Giassi, PhD for histological processing and Manijeh Daneshmand, MD for her histopathological analysis. We thank Virgile Koffi and Nathan Adolphe for pictures in Figure 1A and B.

